# Strategies for the modulation of mitochondrial metabolism and activity in the treatment of neurodegenerative diseases: A systematic review and meta-analysis

**DOI:** 10.64898/2026.03.09.710294

**Authors:** Piedad Valverde-Guillén, Pedro Seoane, Juan A G Ranea, Miguel Ángel Medina, Manuel Marí-Beffa, Beatriz García-Díaz, Manuel Bernal

## Abstract

Neurodegenerative diseases (NDDs) are currently raising their prevalences and new preclinical low-cost investigations of drug design are urging. NDDs encompass a wide range of disorders, including Alzheimer’s, Parkinson’s, ALS and others, many of which share mitochondrial dysfunction as a common pathological feature. As such, targeting mitochondrial metabolism has emerged as a promising therapeutic strategy. However, while rodent models are widely used in NDD research, they are costly and time-consuming, raising the need to consider other alternatives to accelerate the search for novel therapies. In this line, zebrafish (Danio rerio) have gained outstanding popularity as a valuable option. This systematic review aims to provide an extensive overview about the current strategies that use zebrafish assays to investigate modulations of mitochondrial function as new therapies against NDDs. The review was performed following an electronic search of different databases (PubMed, Embase, Scopus and Web of Science) after the PRISMA procedure. Articles published in the English language were identified and screened based on the keywords used: mitochondrial metabolism, therapy, neurodegenerative diseases and zebrafish. Following 176 entries, exclusion criteria reduced the record to 34 final studies. Overall, we found that these studies investigate 37 compounds: 24 natural, 6 semisynthetic, 5 synthetic and 2 compounds of not-determined origin; to ameliorate 9 prevalent diseases: ARSACS, Alzheimer’s, Parkinson’s, Huntington’s diseases, Leigh and Wolfram syndromes, Amyotrophic lateral sclerosis, Limb – girdle muscular dystrophy 2G and hyperglycemia-associated amnesia. Additionally, a meta-analysis of these compounds and their gene interactions provides insights into their mechanisms of action and advances our understanding of NDDs, and furnishes us with a powerful tool to predictive potential new drugs or to repurpose existing ones. To conclude, this systematic review suggests that zebrafish have become an effective model for screening potential drugs for NDDs with symptomatology difficult to replicate in rodent models. Moreover, the use of computational tools is also emphasized as a promising strategy to guide therapeutic discovery more efficiently, reducing both time and costs, in developing treatments for NDDs.

**Graphical abstract:** 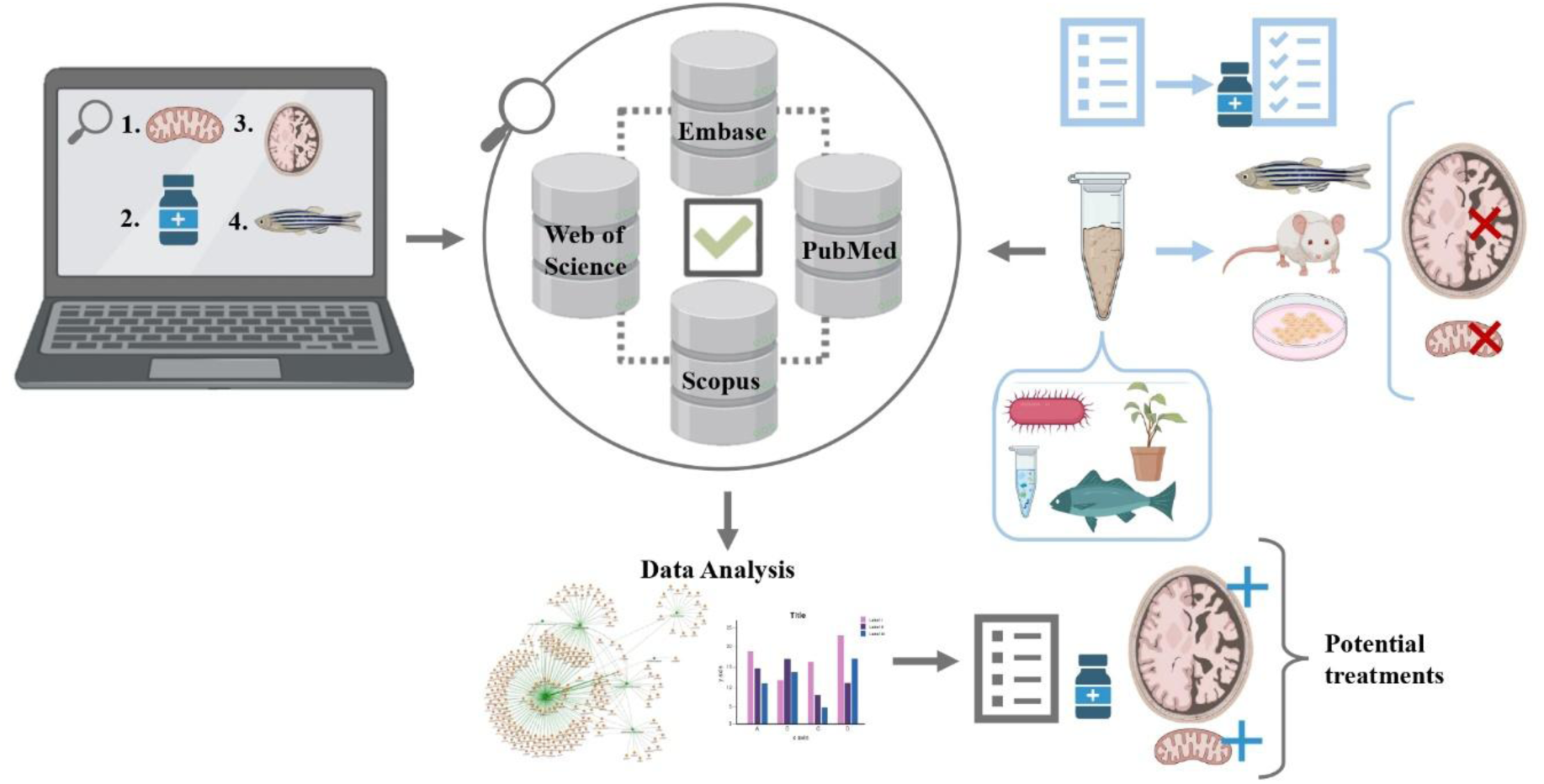

## Introduction

Due to the abrupt aging of the population in the last decades, neurodegenerative diseases (NDDs) are raising their prevalence. This upcoming situation challenges our society to redouble our research efforts in the search for effective treatments to face these escalating health and societal problems. Not surprisingly, finding new treatments for NDDs stands out as the central focus of many research endeavours in recent times. The term NDDs copes with a long list of diseases that includes progressive loss of nerve structure or function containing a broad spectrum of disorders such as amyotrophic lateral sclerosis, multiple sclerosis, Parkinson’s disease, Alzheimer’s disease, Huntington’s disease, multiple system atrophy, tauopathies, and hereditary ataxias, among others. Despite the varied NDD pathomechanisms, mitochondrial dysfunction has been described in many of them as a common feature that links their pathologies. In the last years, many studies on NDD have focused on this aspect, such as in Parkinson’s disease and in other conditions like Leber’s hereditary optic neuropathy^1^, or autosomal dominant optic atrophy ^1^. Thus, the regulation of mitochondrial metabolism and activity has been identified as potentially relevant therapeutic targets for the treatment of NDDs.

The intrinsic characteristics of the nervous system, and the integrative and behavioral outcomes of its dysfunction make almost imperative the use of animal models in the quest for valuable treatments in the study of NDDs. However, the wide use of rodents as animal models for NDDs implies a high time- and economic-cost. As an alternative, zebrafish (*Danio rerio*) has become an established and increasingly popular animal model for the characterization of NDDs and the search for fruitful treatments to defeat them. The relevance of the use of this freshwater fish lays down in the fact that 70% of its genome is shared with humans. Additionally, the zebrafish has a central nervous system organization similar to that of mammals, including humans^2^. Moreover, the ease for gene manipulation allows this species to be used to create quickly and efficiently genetic animal models for specific disease variants. Available gene-editing tools, such as CRISPR-Cas9 variants, enable the modification and breeding of animal models to replicate the specific characteristics of the pathology under investigation, such as Alzheimer’s disease^3^, Parkinson’s disease^4^, among others. In neuroscience, their transparency during the larvae phase allows the visualization of the central nervous system during developmental studies and their social and cognitive abilities identify zebrafish as a good model for behavioral studies. These useful features together with their small size, making easy to breed them in high numbers, and their quick development, which reduces experimental costs and increases research production, spot out zebrafish as an excellent animal model that could be used to generate good platforms for quick and effective testing of drugs.

Highlighting the importance of finding novel strategies to ameliorate or reverse the clinical course of the broad spectrum of NDDs, this systematic review aims to provide a complete overview of the most recent literature about compounds tested in zebrafish animal models with high potential to modulate mitochondrial metabolism and activity that could be tested for therapeutic purposes in the field of NDDs. Moreover, the meta-analysis of the interaction of these reviewed compounds with potential genes and their organization in functional categories provides new perspectives in their therapeutic potential and stands out the importance of using zebrafish as animal model in the study of NDDs.

## Materials and Methods

### Search and Screening Strategy

This review and meta-analysis study were conducted in accordance with the Preferred Reporting Items for Systematic Reviews and Meta-Analysis (PRISMA) 2020 Statement ^5^. A PRISMA checklist is provided in Additional file 1. We conducted a comprehensive literature review, utilizing the following databases: Embase, Web of Science, PubMed, and Scopus in December 2024. Articles published in the English language were evaluated using a specific key term. The search query included keywords combined with Boolean search operators, as follows: mitochondrial metabolism AND therapy AND neurodegenerative diseases AND zebrafish (Table 1). Relevant publications from research and reviews articles published before December 20, 2024, were selected. Studies were screened based on records identified through database searching.

**Table 1.**
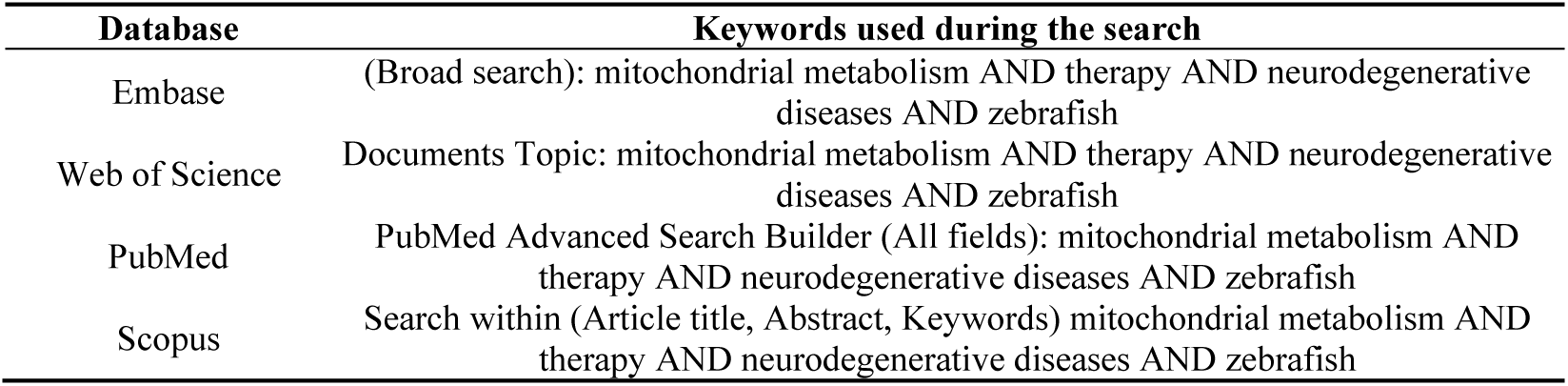
Search terms strategy used during the study.

| Database | Keywords used during the search |
| --- | --- |
| Embase | (Broad search): mitochondrial metabolism AND therapy AND neurodegenerative diseases AND zebrafish |
| Web of Science | Documents Topic: mitochondrial metabolism AND therapy AND neurodegenerative diseases AND zebrafish |
| PubMed | PubMed Advanced Search Builder (All fields): mitochondrial metabolism AND therapy AND neurodegenerative diseases AND zebrafish |
| Scopus | Search within (Article title, Abstract, Keywords) mitochondrial metabolism AND therapy AND neurodegenerative diseases AND zebrafish |

### Eligibility criteria

Articles were screened based on the analysis of the title and abstract according to the predefined inclusion and exclusion criteria (Table 2). If abstracts fit the criteria, a comprehensive full-text analysis was conducted using the same inclusion criteria.

**Table 2.**
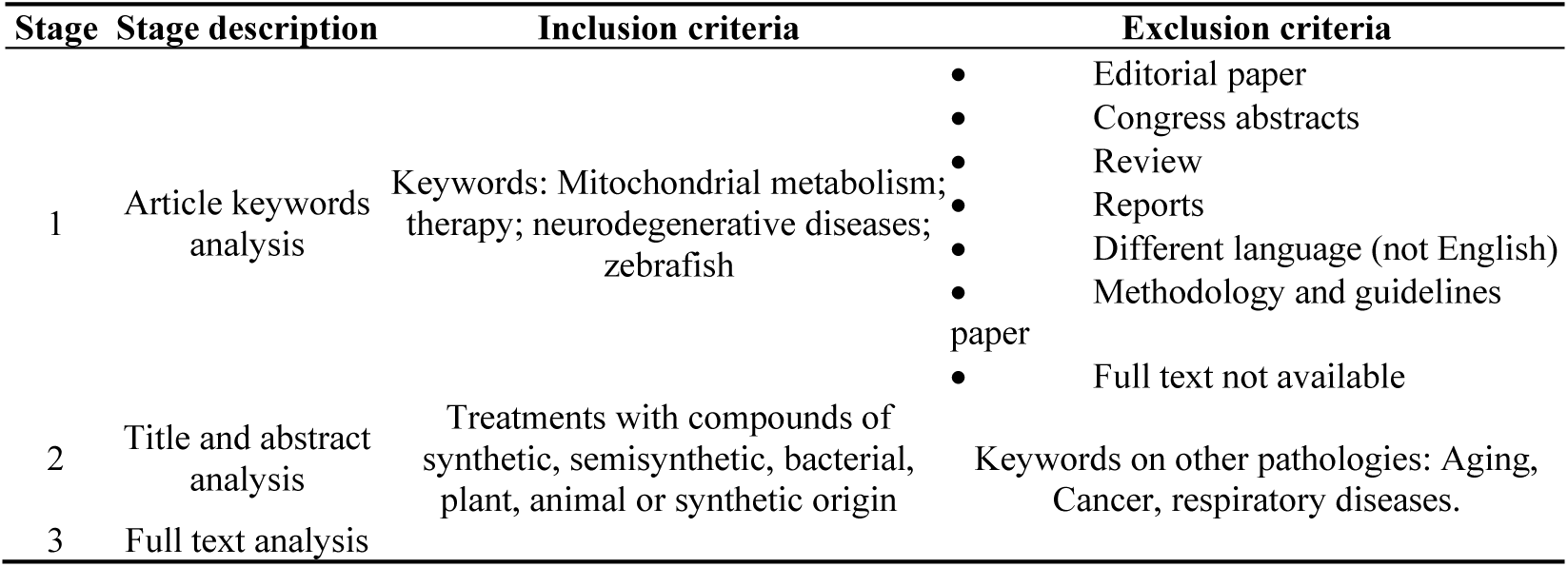
Inclusion and exclusion criteria. At the various stages of analysis for inclusion in the systematic review, papers were included and excluded based on the above criteria.

| Stage | Stage description | Inclusion criteria | Exclusion criteria |
| --- | --- | --- | --- |
| 1 | Article keywords analysis | Keywords: Mitochondrial metabolism; therapy; neurodegenerative diseases; zebrafish | • Editorial paper<br>• Congress abstracts<br>• Review<br>• Reports<br>• Different language (not English)<br>• Methodology and guidelines paper<br>• Full text not available |
| 2 | Title and abstract analysis | Treatments with compounds of synthetic, semisynthetic, bacterial, plant, animal or synthetic origin | Keywords on other pathologies: Aging, Cancer, respiratory diseases. |
| 3 | Full text analysis |  |  |

### Selection of studies

A total of 176 entries were collected from the primary search, using only the databases mentioned above. All references and citations were managed using Mendeley software (Version 2.125.2) to avoid duplication and ensure proper organization ^6^. After removing the duplicate entries, a total of 129 articles remained for the screening of title and abstracts. Among these, 23 entries were eliminated based on the inclusion and exclusion criteria. Subsequently, 106 articles were eligible for full-text assessment, of which 26 studies were excluded due to unavailability of full text. Finally, a total of 34 articles were included for data extraction and analysis in this re-view based on the adopted inclusion criteria. The workflow chart for selection of eligible articles is shown in Figure 1.

**Figure 1.**
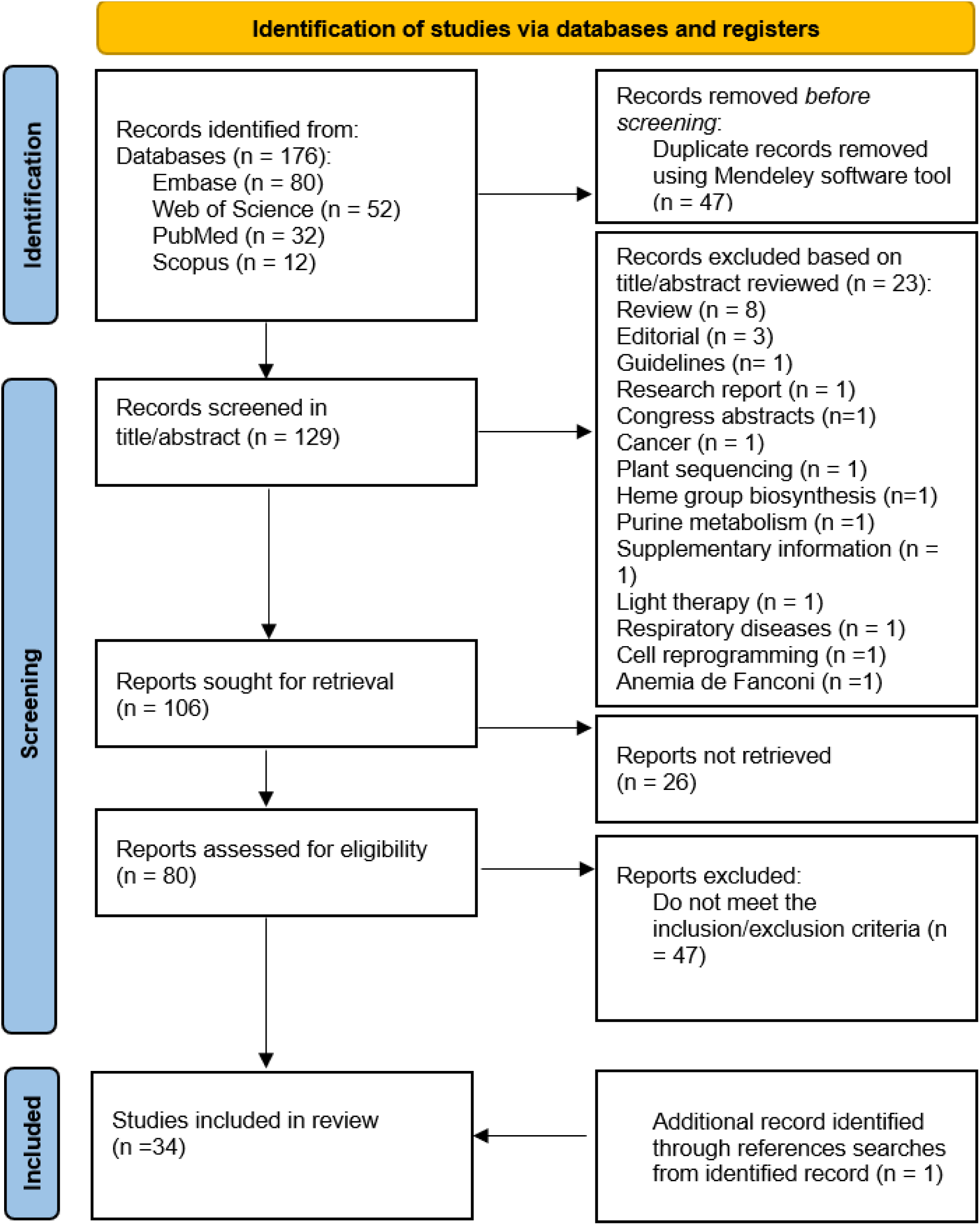
Flow diagram chart of systematic analysis for literature inclusion. Various databases (Embase, Web of Science, PubMed, and Scopus) were utilized to identify all studies published in the English language. We screened 176 articles; 106 were eligible for critical appraisal resulting in a total of 34 articles to be included in this review.

### Data extraction

Two distinct observers, PVG and BGD, performed the literature search independently to identify articles that potentially met the inclusion and identified the reviewed compounds. The extracted data were matched and discussed in a consensus meeting, and disagreements were resolved through discussions between PVG, BGD and MB. Based on the predefined criteria, relevant information about compounds able to modulate mitochondrial metabolism from the eligible studies was extracted and organized in a table by PVG. The extracted information was supplemented using the software tools listed in Table 3.

**Table 3.** Software tools and databases used to supplement the information extracted from the comprehensive analysis of the articles.

| Software tools | Reference |
| --- | --- |
| PubChem | 9 |
| MetaboAnalyst 6.0 | 65 |
| KEGG Database | 66–68 |
| Coconut (COLleCtion of Open Natural ProDUcTs) | 69 |

### Risk of bias and quality assessment

To evaluate the validity of the studies included in this review, two authors (PVG and BGD) independently applied the risk of bias tool developed by the Systematic Review Centre for Laboratory Animal Experimentation (SYRCLE) ^7^. Any discrepancies in assessments were resolved through discussion between the reviewers, or with input from a third independent party (MBM) when necessary.

The SYRCLE tool comprises ten items grouped into six categories: selection bias (sequence generation, baseline characteristics, allocation concealment), performance bias (random housing, blinding), detection bias (random outcome assessment, blinding of outcome assessment), attrition bias (incomplete outcome data), reporting bias (selective outcome reporting), and other potential sources of bias. Each item required a response to a signaling question, classified as “yes” (indicating low risk of bias), “no” (high risk of bias), or “unclear (NC)” (risk of bias not determined). Based on these responses, the risk of bias for each item was categorized as low, high, or unclear. Each included study was assigned a quality score, with a maximum possible score of 10 points.

### Meta-Analysis strategy

Drug-gene interactions were obtained from the online Drug Gene Interaction Database (DGIdb)^8^. Only those with results in the DGIdb^8^ database were included in the meta-analysis. All possible nomenclature variations of each compound were considered based on PubChem^9^. As the database does not allow filtering results by organisms, this process is independent of this criterion.

The compound-disease relationships and the drug-gene interactions from DGIdb were used to build a gene-compound-disease tripartite network. The gene-compound relations were weighted by the interaction score. Evidence scores, the ratio of average known drug partners, and the ratio between average known gene partners for all drugs and the known drug partners for all genes were incorporated into a scoring metric by DGIdb ^8^.

The disease-gene bipartite network was obtained following the methodology of Jabato et al^10^, from the tripartite network, in which the edges represent the number of compounds that connect a disease with a given gene. The network representations were obtained using NetAnalyzer and described in Pagano-Márquez et al^11^.

Gene enrichment meta-analysis was performed for each gene using the genes connected to them in the disease-gene network applying the methodology for functional analysis described in detail by Pagano-Márquez et al^11^.

## Results

### Molecules as potential regulators of mitochondial metabolism for treating neurodegenerative diseases

We identified 176 records from our literature search. After screening, 106 were eligible for critical appraisal resulting in a total of 34 articles to be included in this systematic review (Figure 1). Once extracted the list of the used compounds able to modulate mitochondrial in the reviewed articles, their characteristics (chemical nature, chemical source and involved physiological pathways) were completed using the software tools (Table 3) and organized in a table (Table 4) to provide a whole overview of the molecules with potential capacities to treat NDDs via targeting mitochondrial metabolism.

**Table 4.**
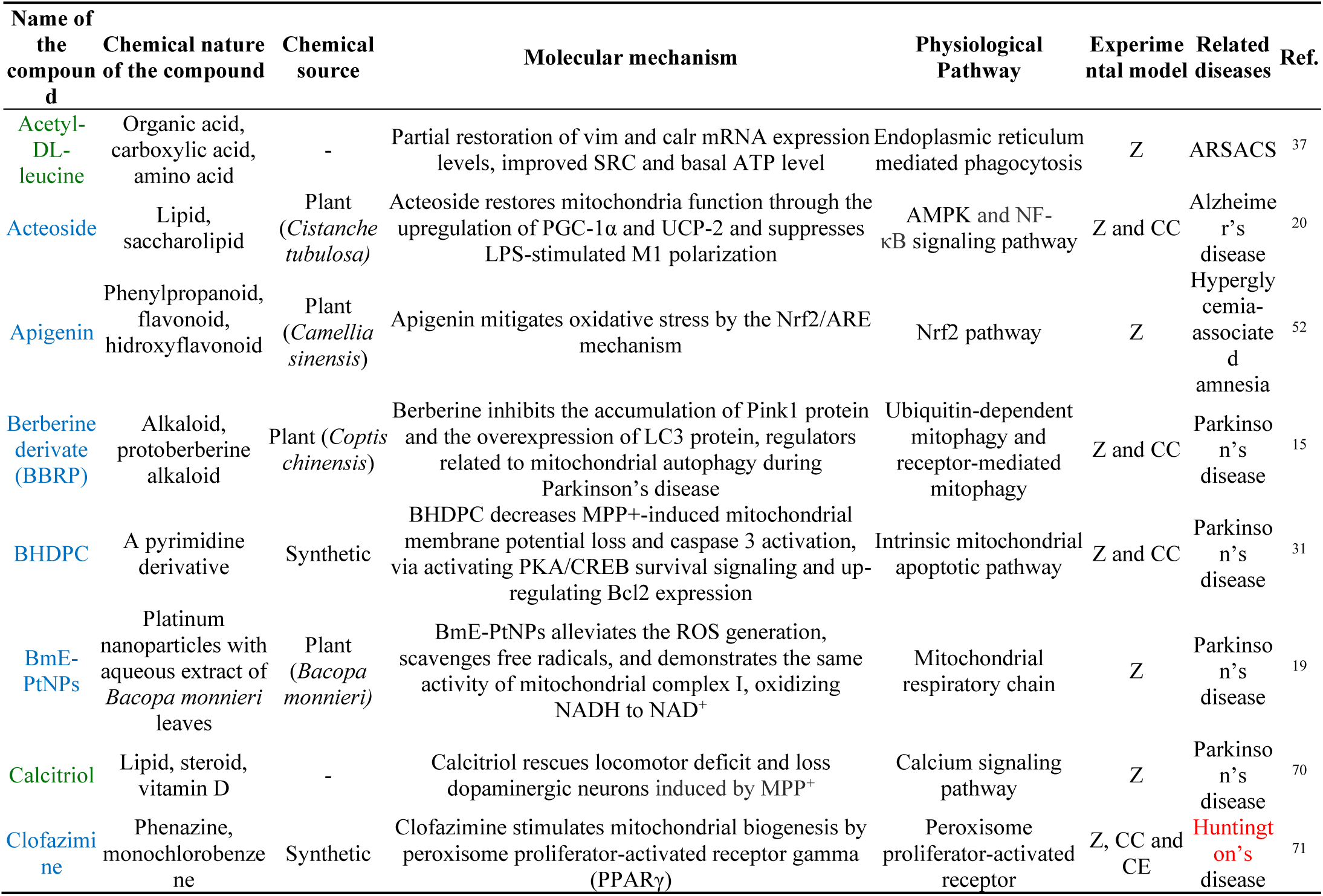
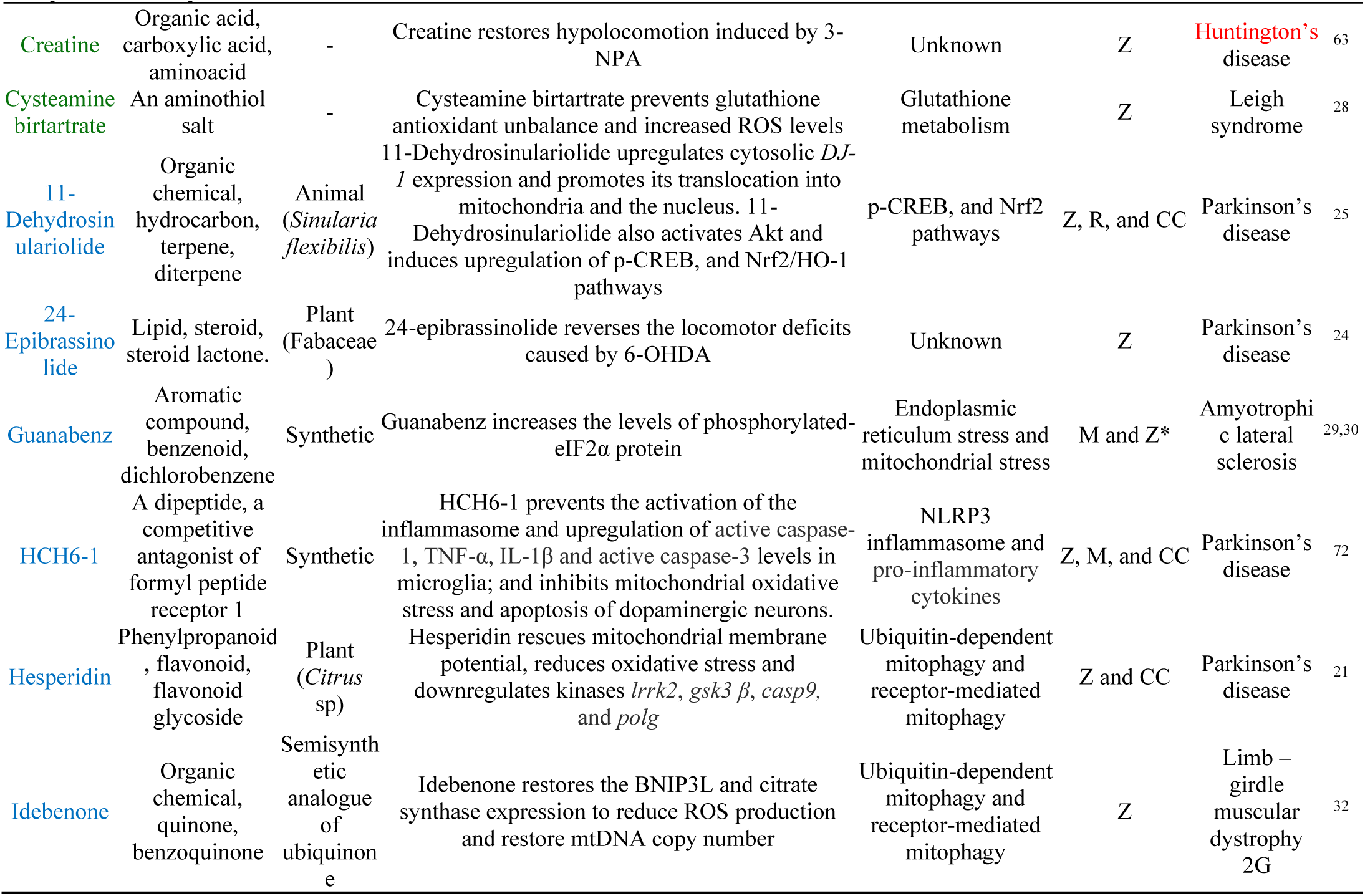
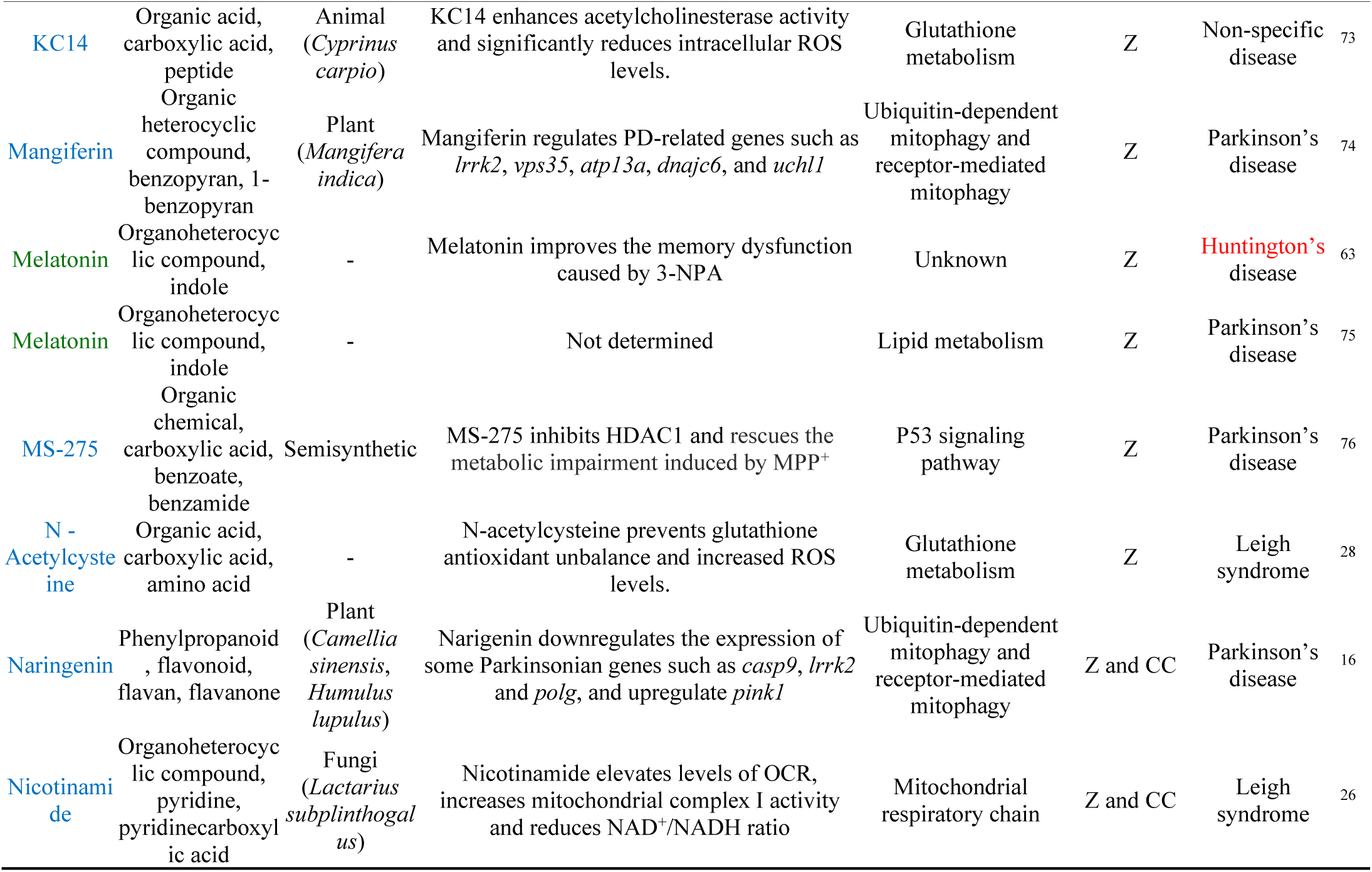
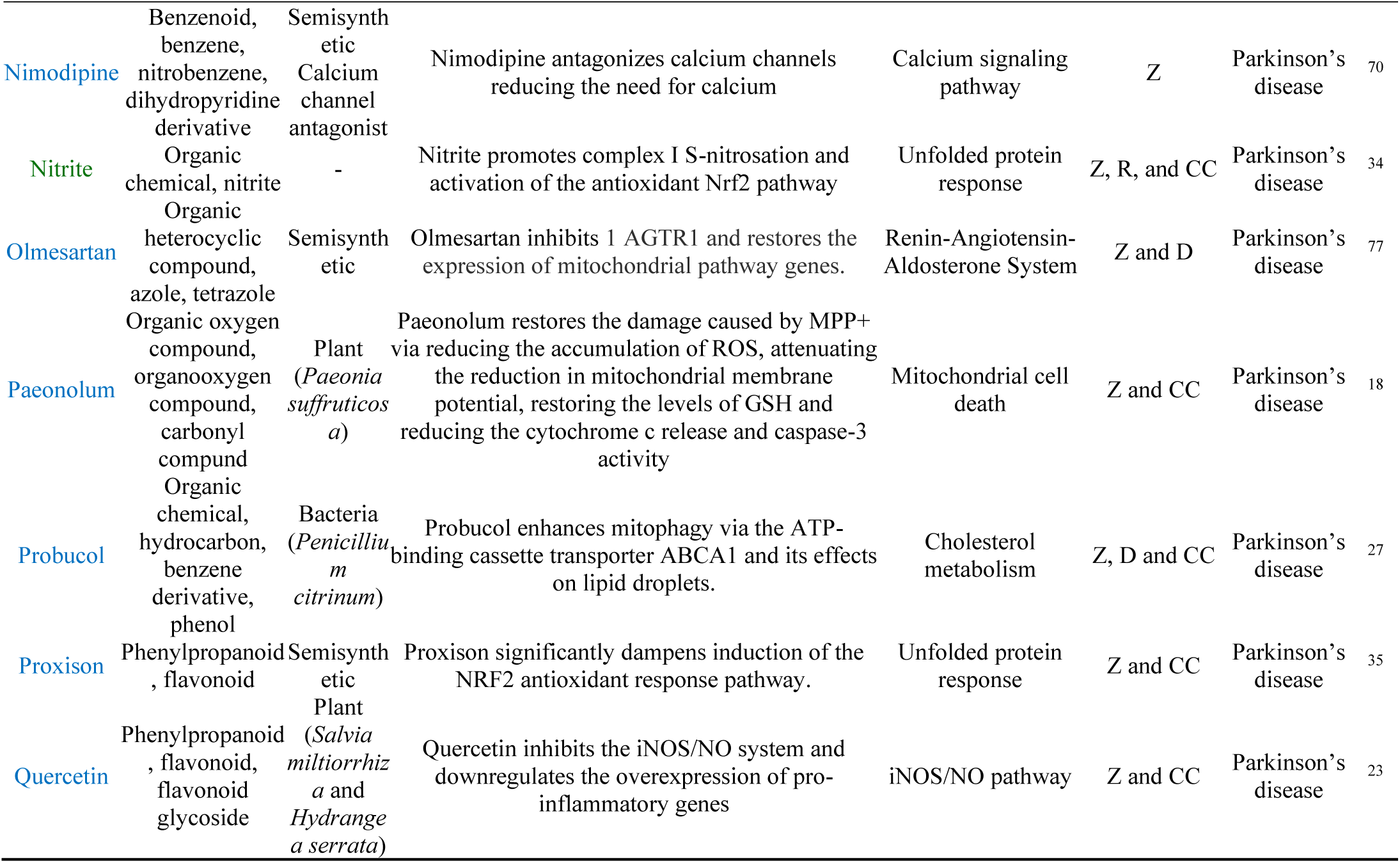
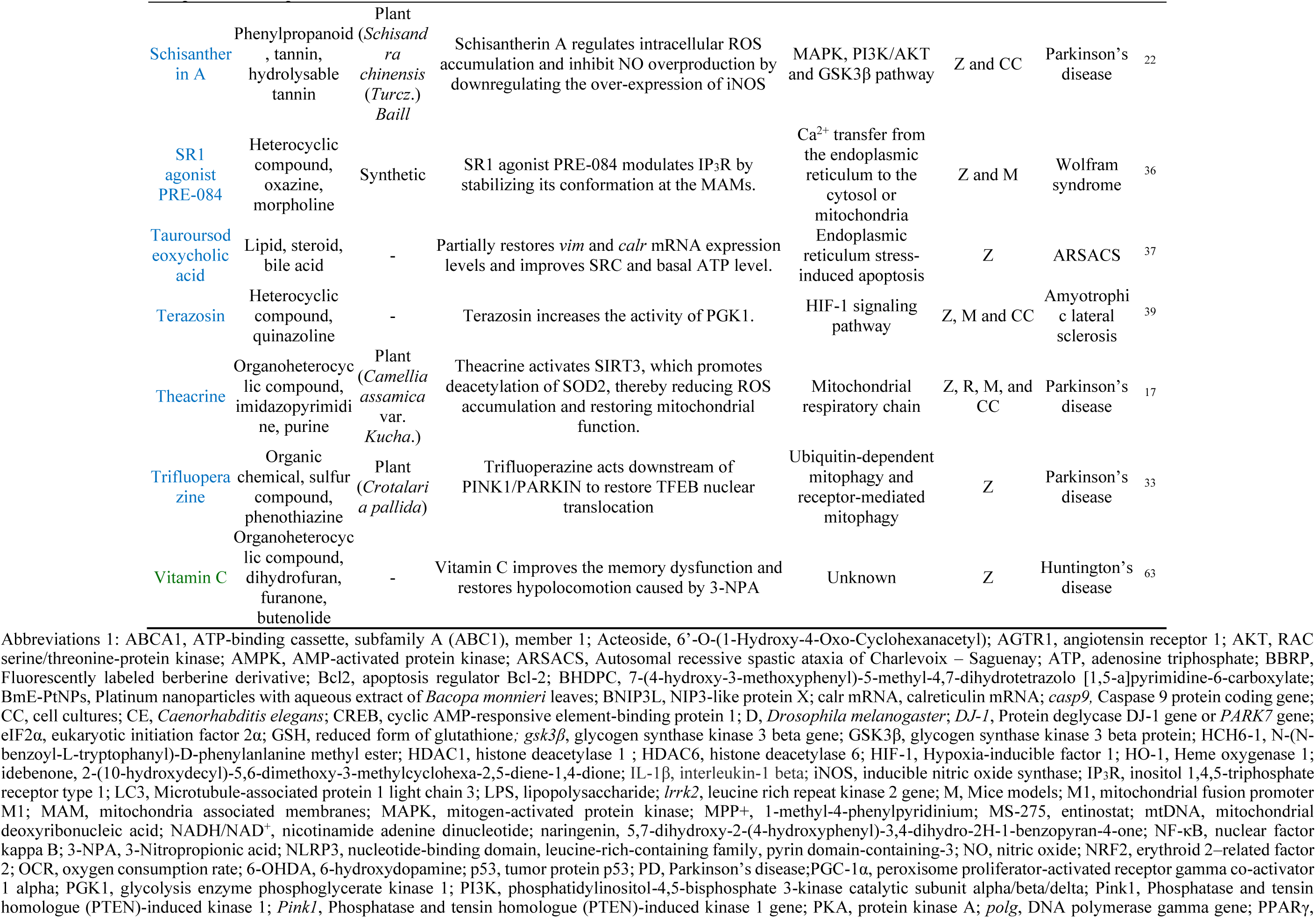
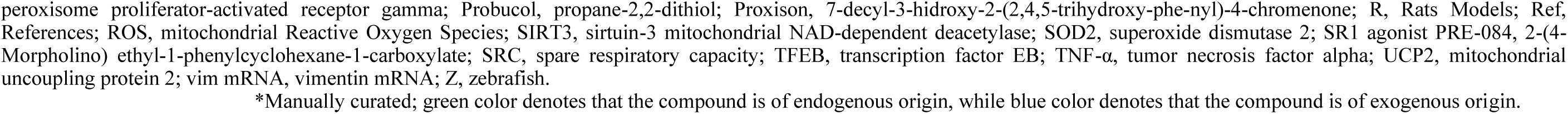
Compounds of endogenous or exogenous origin and their known molecular characteristics in the treatment of neurodegenerative diseases.

### Methological Quality Assessment

In recent times, studies that propose new treatments to alleviate various NDDs have focused on seeking modulators of mitochondrial metabolism, which is dysfunctional in many of these pathologies^12^. Animal experimentation remains crucial for evaluating drugs in the context of neurodegenerative diseases. However, aiming to estimate the significance of the assumptions gathered from animal studies, methodological quality should be evaluated. To assess the methodological quality of the studies included in this systematic review, the SYRCLE tool has been employed and scored data has been summarized in table 5.

**Table 5.** Methodological quality using SYRCLE tool.

|  | 1 | 2 | 3 | 4 | 5 | 6 | 7 | 8 | 9 | 10 | Score |
| --- | --- | --- | --- | --- | --- | --- | --- | --- | --- | --- | --- |
| Zhang ZJ et al. (2011)<sup>23</sup> | Y | Y | NC | NC | NC | NC | NC | NC | Y | Y | 4 |
| Nellore J et al. (2013)<sup>19</sup> | NC | NC | NC | NC | NC | NC | NC | NC | Y | Y | 2 |
| Vaccaro A et al. (2013)<sup>30</sup> | NC | NC | NC | NC | NC | NC | NC | NC | Y | NC | 1 |
| Jiang HQ et al. (2014)<sup>29</sup> | Y | Y | NC | NC | NC | NC | Y | NC | Y | NC | 4 |
| Chong C et al. (2015)<sup>31</sup> | NC | Y | NC | NC | NC | NC | NC | NC | Y | Y | 3 |
| Lu XL et al. (2015)<sup>18</sup> | NC | Y | NC | NC | NC | NC | Y | NC | Y | Y | 4 |
| Pinho BR et al. (2015)<sup>76</sup> | Y | Y | NC | NC | NC | NC | NC | NC | Y | Y | 4 |
| Zhang LQ et al. (2015)<sup>22</sup> | NC | Y | NC | NC | Y | NC | Y | NC | Y | NC | 4 |
| Feng CW et al. (2016)<sup>25</sup> | Y | Y | NC | NC | Y | NC | Y | NC | Y | NC | 5 |
| Drummonf NJ et al. (2017)<sup>35</sup> | NC | Y | NC | NC | NC | NC | NC | NC | Y | NC | 2 |
| Zhang Y et al. (2017)<sup>33</sup> | NC | Y | NC | NC | NC | NC | NC | NC | Y | Y | 3 |
| Milanese C et al. (2018)<sup>34</sup> | NC | Y | NC | NC | NC | NC | Y | NC | Y | NC | 3 |
| Duan WJ et al. (2020)<sup>17</sup> | Y | Y | NC | NC | NC | NC | NC | NC | Y | NC | 3 |
| Kesh S et al. (2021)<sup>21</sup> | NC | Y | NC | Y | NC | NC | NC | NC | Y | Y | 4 |
| Kesh S et al. (2021)<sup>16</sup> | NC | Y | NC | NC | NC | NC | NC | NC | Y | Y | 3 |
| Kim GHJ et al. (2021)<sup>77</sup> | NC | Y | NC | NC | Y | NC | Y | NC | Y | Y | 5 |
| Li Y et al. (2021)<sup>20</sup> | NC | Y | NC | NC | NC | NC | NC | NC | Y | NC | 2 |
| Naef V et al. (2021)<sup>37</sup> | Y | Y | NC | NC | NC | NC | NC | NC | Y | Y | 4 |
| Wang L et al. (2021)<sup>15</sup> | NC | Y | NC | NC | NC | NC | NC | NC | Y | NC | 2 |
| Chaytow H et al. (2022)<sup>39</sup> | Y | Y | NC | NC | Y | NC | Y | NC | Y | Y | 6 |
| Crouzier L et al. (2022)<sup>36</sup> | NC | Y | NC | Y | Y | NC | Y | Y | Y | Y | 7 |
| Lv X et al. (2022)<sup>32</sup> | NC | Y | NC | NC | NC | Y | NC | NC | Y | NC | 3 |
| Gomes A et al. (2023)<sup>24</sup> | Y | Y | NC | NC | NC | NC | NC | Y | Y | Y | 5 |
| Haroon S et al. (2023)<sup>28</sup> | NC | Y | NC | NC | NC | NC | NC | NC | Y | NC | 2 |
| Moskal N et al. (2023)<sup>27</sup> | NC | Y | NC | NC | NC | NC | NC | NC | Y | NC | 2 |
| Wang HL et al. (2023)<sup>72</sup> | NC | NC | NC | NC | NC | NC | NC | NC | Y | NC | 1 |
| Haridevamuthu B et al. (2024)<sup>52</sup> | Y | Y | NC | NC | NC | Y | Y | NC | Y | Y | 6 |
| Kim M et al. (2024)<sup>70</sup> | NC | Y | NC | NC | NC | NC | NC | NC | Y | Y | 3 |
| Li X et al. (2024)<sup>71</sup> | Y | Y | NC | NC | NC | NC | NC | NC | Y | NC | 3 |
| Lo CH et al. (2024)<sup>26</sup> | NC | Y | NC | NC | NC | NC | NC | NC | Y | NC | 2 |
| Pang MZ et al. (2024)<sup>75</sup> | NC | Y | NC | NC | NC | NC | NC | NC | Y | NC | 2 |
| Qin F et al. (2024)<sup>74</sup> | Y | Y | NC | NC | NC | NC | NC | NC | Y | NC | 3 |
| Vijayanand M et al. (2024)<sup>73</sup> | NC | Y | NC | NC | NC | NC | NC | NC | Y | NC | 2 |
| Wiprich M et al. (2024)<sup>63</sup> | Y | Y | NC | NC | NC | NC | NC | NC | Y | NC | 3 |

The quality items score of each study is ranged from one to seven points, with a mean of 3.29 out of a total 10 points. In these studies, the main risk of bias lies on the lack of information (NC). No study declares a proper concealed allocation, and only one (2,94%) states random housing. Although the risk of bias in these studies due to the lack of randomization might be medium-low (items 4 and 6), the majority of the articles acknowledge very similar baseline characteristics (item 2 “Y” in 31, 91.17%), and proper reporting bias (item 9 “Y” in 34, 100%) free of selective outcome reports.

However, there is a main type of high risk of bias among these publications that is the lack of blinding both in caregivers and investigators (item 5 “Y” only in 5 studies, 14,70%) and in the outcome assessors (item 7 “Y” only in 9 articles, 26,47%). In addition, few publications mention sample size calculation. The sample size in animal studies should be large enough to detect biologically significant differences, while also being small enough to minimize unnecessary animal sacrifice. Therefore, this lack of information could harm the internal validity of the evidence from these animal studies.

### Analysis of Potential Mitochondrial Regulator Characteristics

In line with the search for drugs with potential therapeutic capacities via regulating mitochondrial metabolism to improve, prevent, or even cure the mentioned pathologies, numerous studies focus on testing compounds of diverse origins. In our reviewed list, some compounds could be considered of endogenous origin (green color in table 4), as they are naturally produced by the body, such as neuroglobin^13^ and coenzyme Q10 ^14^ (Metabolite (23%) in Figure 2), while the majority have an exogenous origin (blue color in table 4) (Plant (33%), Animal (5%), Bacteria (3%), Synthetic (13%), Semisynthetic (15%), Fungi (3%), and Unknown (total 77%) in Figure 2). Among those latter, most of the studies that presented a mitochondrial regulator compound of exogenous origin, i.e. Berberine derivative^15^, naringenin^16^, theacrine^17^, paeonolum^18^, BmE-PtNPs^19^, acteoside^20^, hesperidin^21^, schisantherin A^22^, quercetin^23^, 24-epibrassinolide^24^, and 11-dehydrossinulariolide^25^ have a naturally occurring chemical origin, as they are extracted from various plant species (33%), being compounds of fungi^26^, animal^25^, bacterial^27^ origin and those with no precise mention of their origin in the study the least represented (Figure 2).

**Figure 2.**
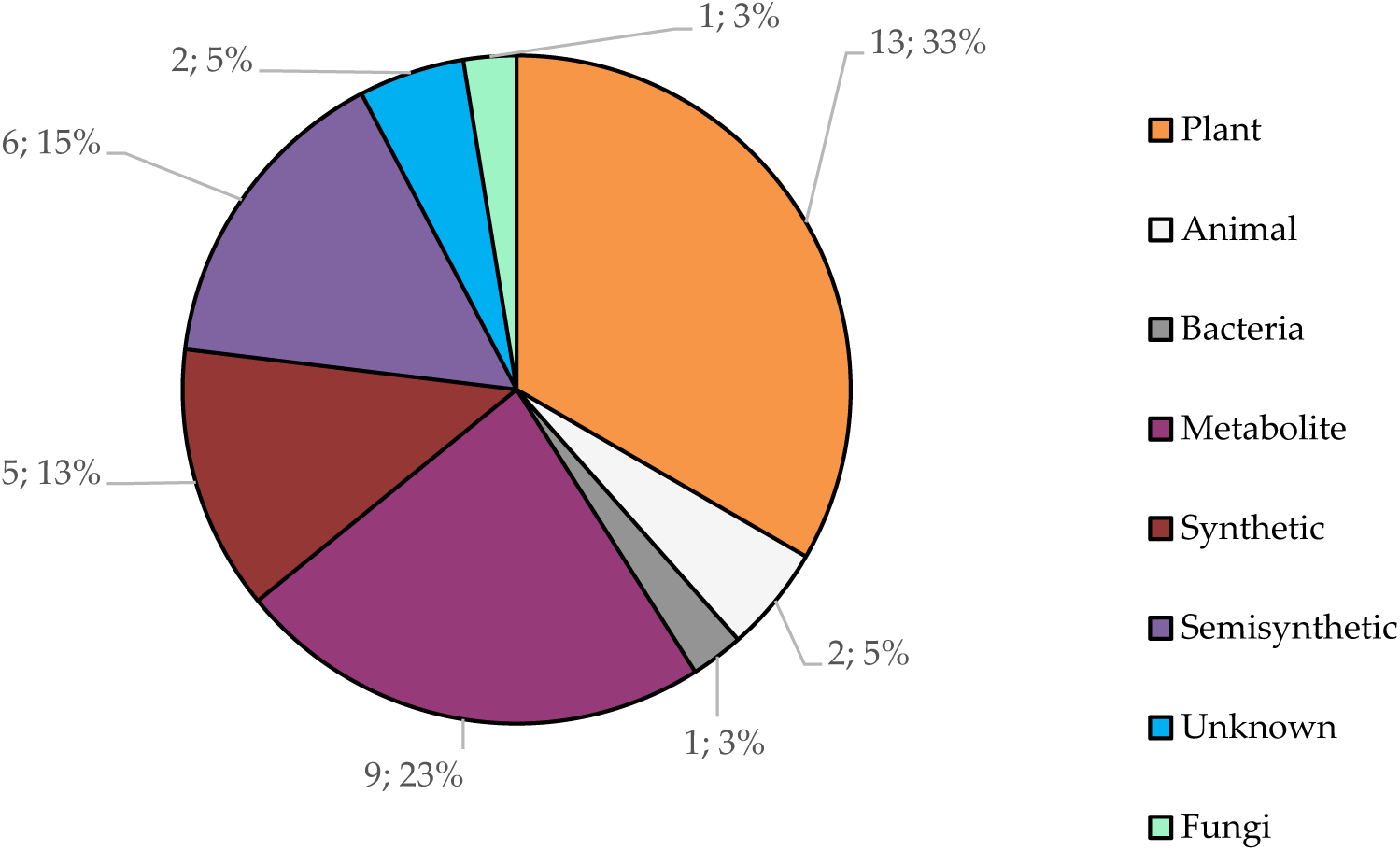
Pie chart representing the number of compounds discussed in this review by their chemical source.

### Mitochondrial Regulation Mechanisms Analysis

Regarding the mode of action of the studied compounds, most of them affect directly molecular mechanisms related to mitochondrial metabolism and their activity per se, such as mitochondrial respiratory chain^17,19,28^, mitochondrial stress^29,30^, mitochondrial cell death^18^, intrinsic mitochondrial apoptotic pathway^31^, and ubiquitin-dependent mitophagy or receptor-mediated mitophagy^15,16,32,33^, being mitophagy the mechanism more frequently targeted by the reviewed drugs.

Whereas, other compounds also involve mode of actions that regulate the activities of other organelles or cell functions, likely upstream to mitochondrial function or downstream consequences of mitochondrial dysfunctions, such as endoplasmic reticulum^34,35^, i.e. Ca^2+^ transfer from the endoplasmic reticulum to the cytosol or mitochondria^36^, endoplasmic reticulum stress-induced apoptosis^29,30^, endoplasmic reticulum mediated phagocytosis ^37^, or cell signalling pathways, such as AMPK and NF-κB^20^, p-CREB and Nrf2^25^, P53^38^, iNOS/NO^23^, MAPK, PI3K/AKT and GSK3β^22^, or HIF-1 signalling pathways^39^ (Table 4; Figure 3).

**Figure 3.**
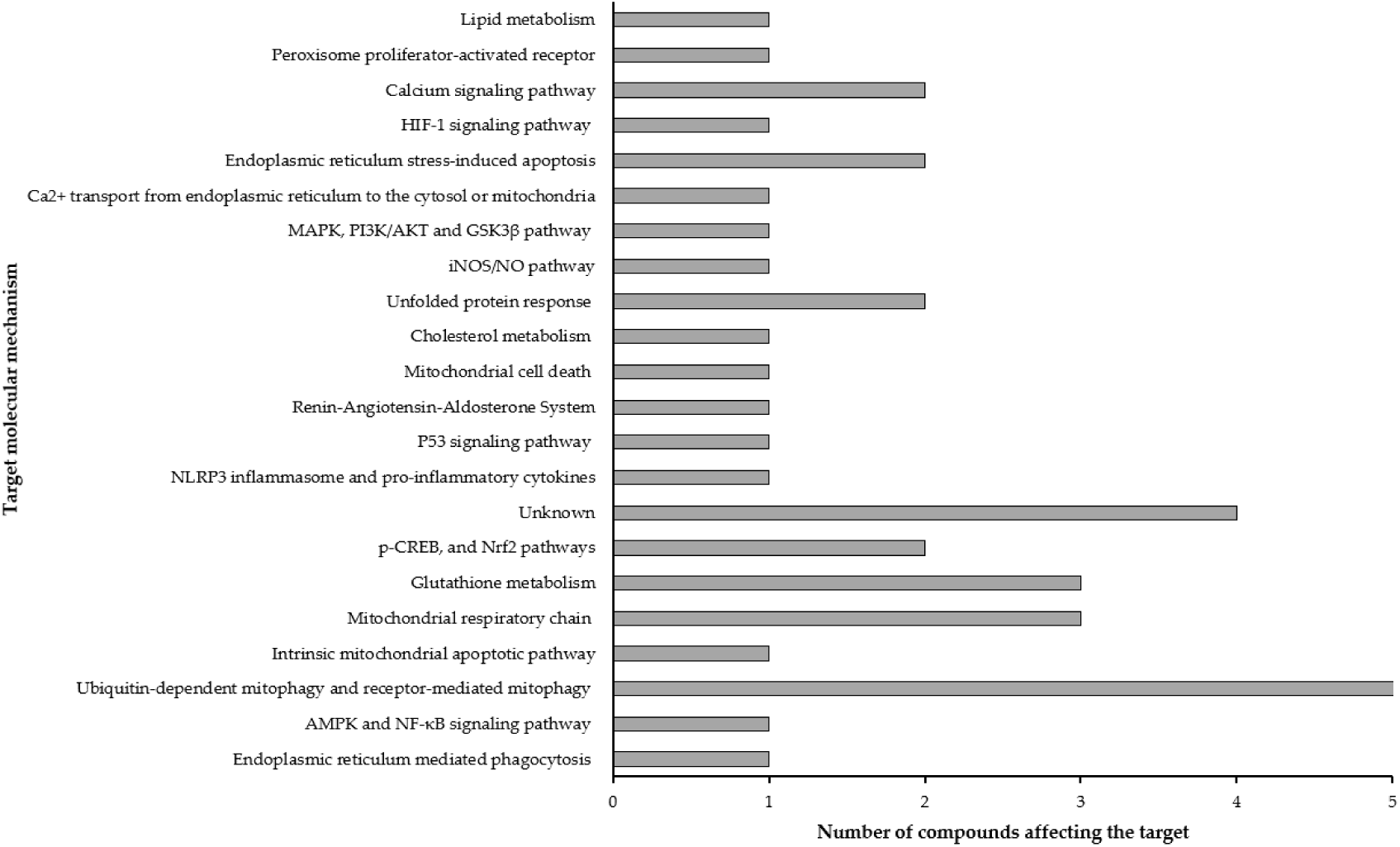
Bar chart representing the total number of molecular mechanisms affected by compounds reviewed.

### Analysis of the Neurodegenerative Disease Models

An interesting aspect is the high number of articles that search for therapies against Parkinson’s disease in zebrafish, testing 22 out 37 different compounds. The other investigated NDDs have drawn less attention, being followed by far for Huntingtońs disease (4 out 37) Leigh syndrome (3 out 37) and Amyotrophic lateral sclerosis (2 out of 37), while the rest of the disease are tackled by just one study and/or one compound (Figure 4).

**Figure 4.**
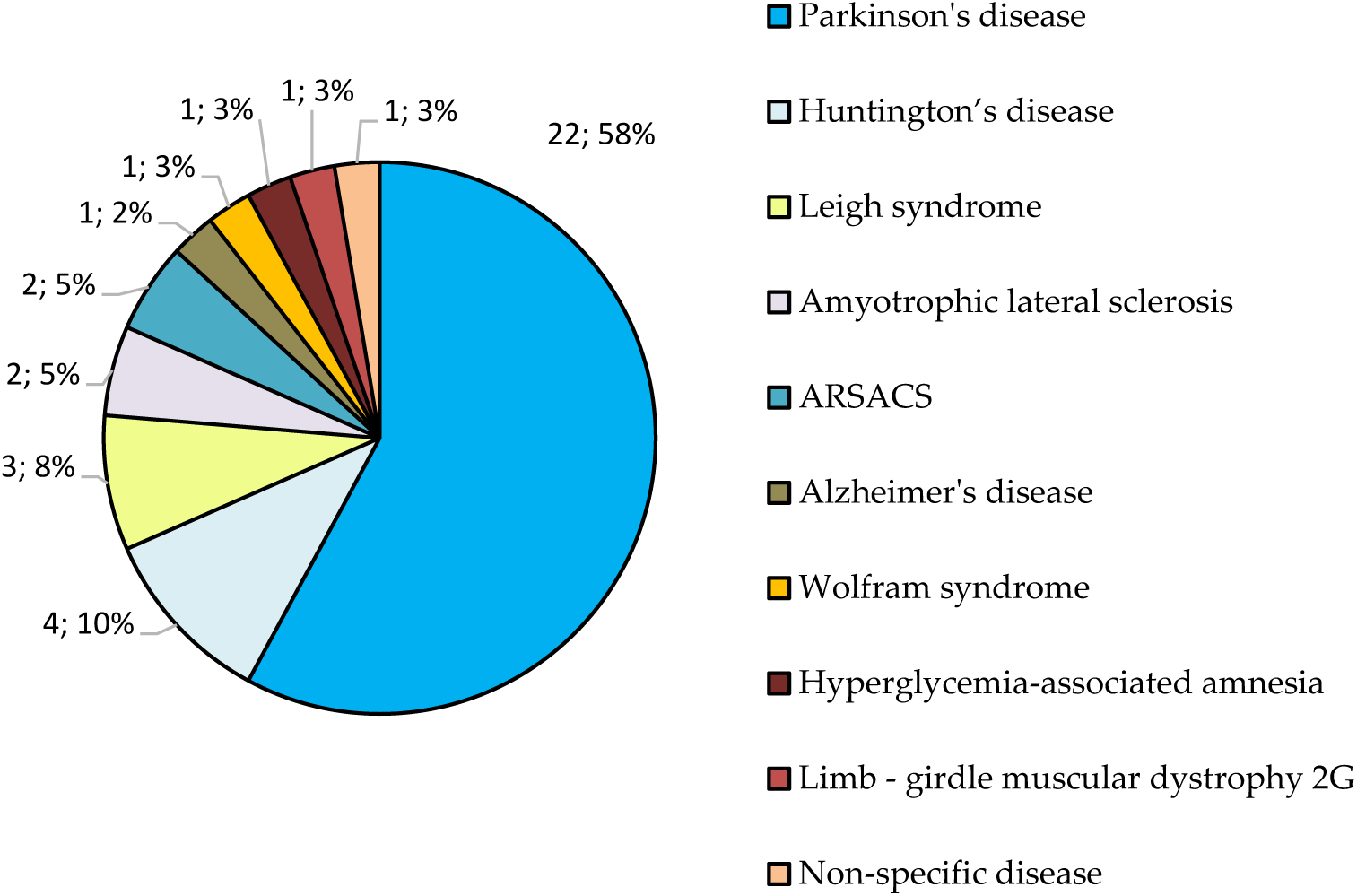
Representation of the number of neurodegenerative diseases targeted by the compounds discussed in this review.

Parkinson’s disease is the second most prevalent NDD, with progressive depletion of dopaminergic neurons in substantia nigra pars compacta ^40^. The cause of the disease is unknown, but there are some potential risk factors, such as genetic factors^41^, age^42^ or environmental agents^43^ that must be taken into account. Although no better animal models are available for the other mentioned NDDs in this study, the notable prevalence of Parkinson’s disease, as identified through this systematic search, is an aspect that warrants further analysis.

### Meta-Analysis of Drug-Gene Interactions and functional enrichment

We conducted a study on the interactions between the selected compounds used as potential treatments for various NDDs (Table 4) and their known target genes. Of the compounds listed in Table 4, only those with results in the DGIdb^8^ database were included in the meta-analysis (Table S1). All possible nomenclature variations of each compound were considered based on PubChem^9^. As the database does not allow filtering results by organisms, this process is independent of this criterion.

Then, we collected drug-gene interactions from the online DGIdb for the curated list of compound-disease relationships generated in this study. The compound-disease relationships and the drug-gene interactions from DGIdb were used to build a gene-compound-disease tripartite network. The compound disease relationships are binary connections that only represent the evidence of the disease affectation by the compound. In the case of gene-compound relations, they are weighted by the interaction score. The DGIdb calculates a scoring metric that incorporates evidence scores, the ratio of average known drug partners, and the ratio between average known gene partners for all drugs and the known drug partners for all genes ^8^. From this tripartite network (Figure 5), we obtained a disease-gene bipartite network (Figure 6) in which the edges represent the number of compounds that connect a disease with a given gene.

**Figure 5.**
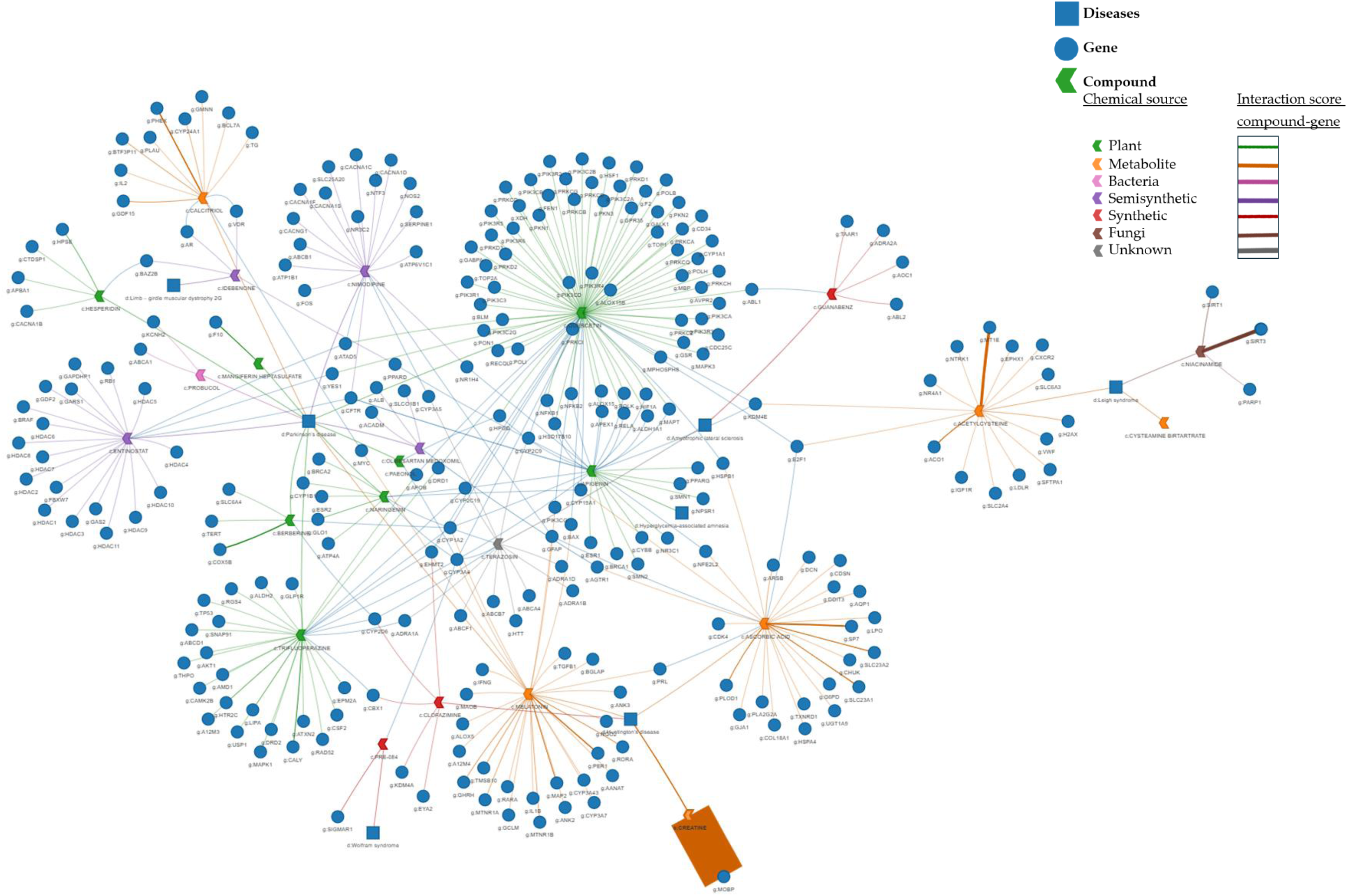
Network of drug-gene interactions in neurodegenerative diseases. Interaction network of the drugs identified through the systematic search and genes obtained from the DGIdb database related to neurodegenerative diseases. Blue circular nodes represent genes. Arrow nodes represent compounds, with their color indicating the chemical source of the compound (green, plant origin; orange, metabolite; pink, bacteria; purple, semisynthetic; red, synthetic; brown, fungi and grey, unknown). Blue square nodes represent the diseases associated with the drugs. The lines represent drug-gene interactions, with thickness of the line indicating interaction scores.

**Figure 6.**
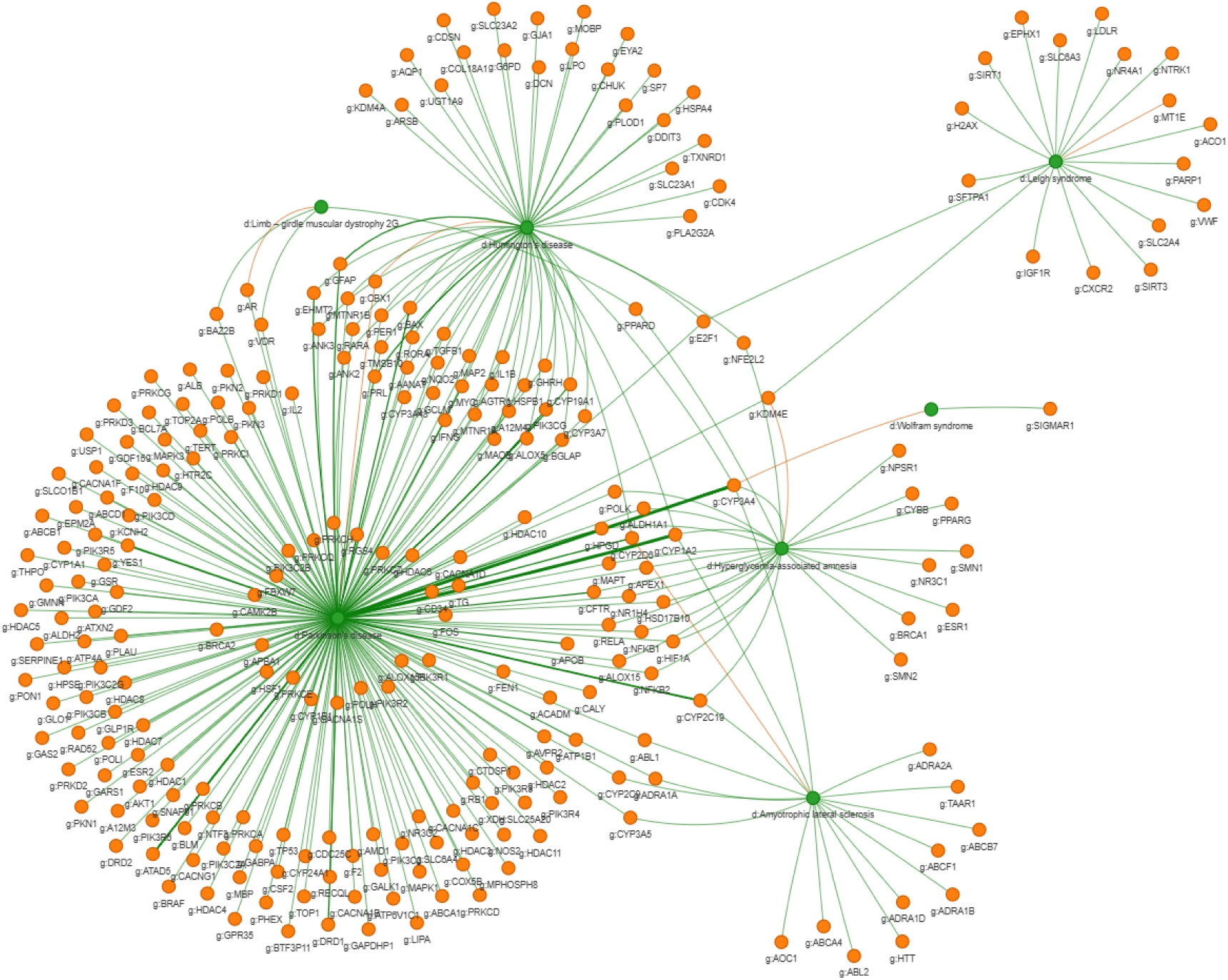
Projected network of gene interactions in relation to neurodegenerative diseases. Interaction network of genes related to drugs obtained from the DGIdb and diseases identified through a systematic review search. Green circular nodes represent diseases. Orange circular nodes represent the genes. Lines represent disease-gene relationships, with line thickness indicating the inferred interaction scores.

As shown in Figure 5, the tripartite network displays some disease nodes with a higher number of known connections to genes. Notably, plant-derived compounds interact with more common genes, as observed in the network’s dispersion. The most connected genes in the network are *CYP3A4*, *CYP1A2*, *CYP2C19*, *CYP2D6*, *ATAD5*, *CFTR*, *CYP19A1*, *E2F1*, *EHMT2* and *GFAP*.

Examining the interaction strength between the compound and the gene, the case of the compound creatine and the gene *MOBP* is particularly striking, as it shows a much stronger interaction compared to the other interactions between the nodes (Figure 5).

A gene enrichment meta-analysis (Figures 7 and 8) was finally performed for each gene using the genes connected to them in the disease-gene network applying the methodology for functional analysis described in detail by Pagano-Márquez et al^11^.

**Figure 7.**
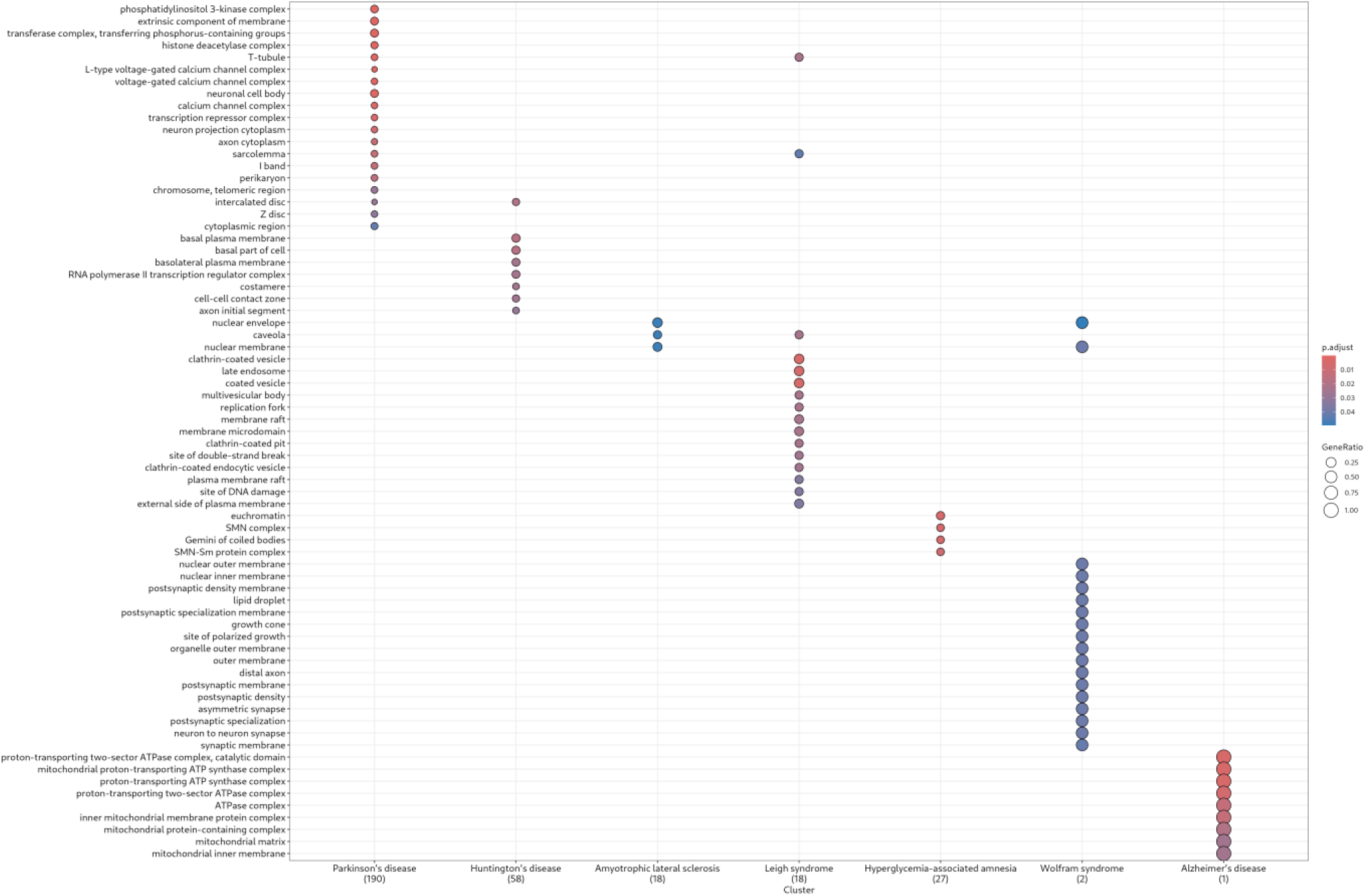
Dot plot of biological pathway categories obtained using the gene ratio, defined as the proportion of significant genes found in each functional category. The x-axis represents the diseases with the gene ratio, and the dot size indicates the number of genes associated with each functional category.

**Figure 8.**
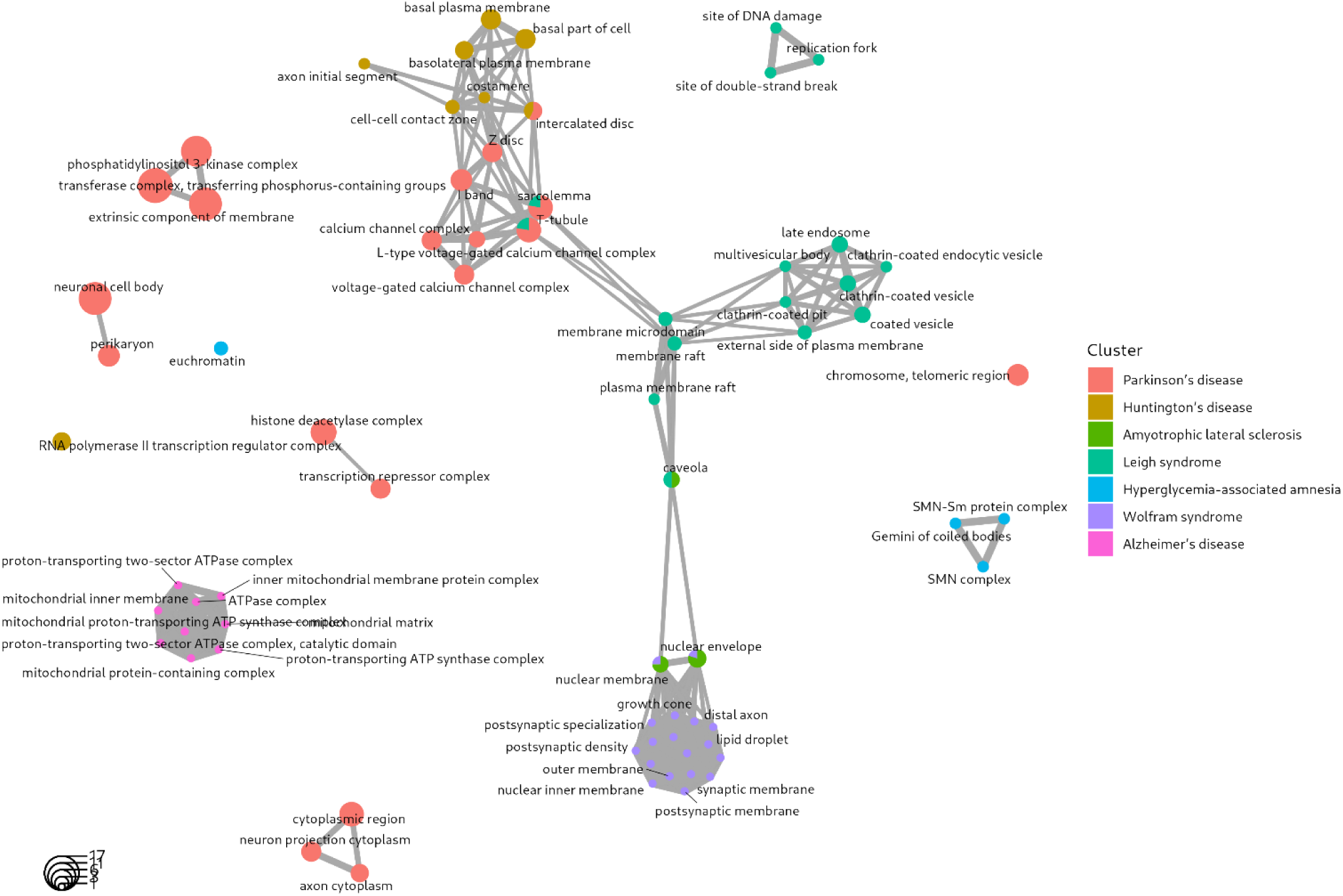
Functional analysis of neurodegenerative diseases. Enrich map plot of cellular location categories obtained using the gene ratio, defined as the proportion of significant genes found in each functional category. The top functional cellular location categories (nodes) are connected if they share genes. Edge thickness represents the number of shared genes and node size represents the number of significant genes within each category. Pie chart proportions represent the proportion of genes belonging to each cluster.

Regarding the molecular elements and pathways involved in the pathologies included in the meta-analysis (Parkinson’s disease, Huntington’s disease, Amyotrophic lateral sclerosis, Leigh syndrome, Hyperglicemia-associated amnesia, Wolfram syndrome and Alzheimer’s disease), we can observe the results in Figure 7, based on the network projection of genes interacting with the compounds and the diseases associated with each study (Figure 6).

It is noteworthy to highlight the dispersion of results, where each disease presents, almost without overlap, a specific set of affected cellular elements. In the context of Parkinson’s disease, various types of calcium-dependent channels are identified, with the genes encoding them interacting with the compounds tested in this disease’s context. Moving on to Huntington’s disease, the most relevant elements are associated with membrane regions, basal membrane, basolateral membrane, or segments related to these elements. Amyotrophic lateral sclerosis is more closely linked to therapeutic targets in caveolae and nuclear membranes. Leigh syndrome is associated with membrane regions involved in endocytosis and exocytosis processes, such as endosomes and clathrin-associated vesicles. In hyperglycemia-associated amnesia, the relevant regions are those of the nucleus, including both the inner and outer membranes. In Wolfram syndrome, specific regions of the postsynaptic and presynaptic membranes in the neuronal context appear to be implicated. In contrast, in Alzheimer’s disease, ATPases of various natures seem to be of relevance (Figure 7).

In Figure 8, we can also observe the dispersion of each point representing the regions seemingly affected in each disease. It becomes apparent that the involvement of specific regions and pathways in the cell, related to the disease pathogenesis, could lead to the identification of specific drugs whose targets are the modulation of certain differential genes in each pathology (Figures 5 and 6). Thus, in Figure 6, the fact that some genes are only associated with one disease may suggest that they represent a potential target for the specific disease.

## Discussion

Zebrafish have emerged as a valuable and cost-effective animal model for studying potential therapies targeting mitochondrial metabolism in NDDs. Their unique biological features—such as transparent embryos, rapid development, and approximately 70% genetic homology with humans—make them particularly suited for *in vivo* studies that require high-throughput screening and visualization of neural and mitochondrial processes.

In the studies reviewed, the majority of compounds tested for their neuroprotective effects were of natural origin, primarily derived from plants. These bioactive molecules act through diverse mechanisms centred on mitochondrial regulation, including boosting ATP production, reducing oxidative stress, stimulating mitophagy, and suppressing apoptotic signalling.

Interestingly, a significant portion of the zebrafish-based NDD research has focused on Parkinson’s disease. Although the reason for that remains speculative, one potential explanation could be that genetic models in rodents have not fully replicated the clinical and neurological characteristics of the pathology^41^, as age-dependence and progressive symptoms. In efforts to generate new disease models, chemicals have often been used as an alternative, which have been described to reproduce the symptoms of the disease^43^. One of these chemicals compounds is 6-OHDA. This synthetic compound, analogous to norepinephrine and dopamine^44^, is used to induce Parkinsonian symptoms in animal models. However, 6-OHDA does not cross the blood-brain barrier (BBB), therefore is needed to be injected into the brain of rodents to create a lesion, while only be exposed to it in zebrafish ^16^. Neurotoxins with the same effect that does cross the BBB, including MPP+^31^, are also found, but resulting in different mechanism of action^45^.

Zebrafish assays to test drugs against human diseases are becoming popular during recent years^46–49^. Among these assays, only a few succeeded in their purpose, being intensively used in drug screenings against osteoporosis^46–49^. In these cases, zebrafish assays were found even more useful than rodent assays for specific metabolic reasons, such as osteoporosis inducibility by anti-inflammatory treatments, a peculiar response in humans not found in mice. Regarding NDDs, zebrafish opens up new possibilities to be used for high-throughput drug screening, while the meta-analysis of the information obtained from the tested compounds can provide interesting clues for the developing and repurposing of drug in the context of NDDs as shown by our results.

*CYP3A4,* one of the most connected genes in the gene-compound-disease tripartite network, encodes in humans a member of the cytochrome P450 superfamily of enzymes. The cytochrome P450 superfamily is a group of monooxygenase enzymes involved in drug metabolism and the biosynthesis of cholesterol and other lipids. This molecular pathway has a relevant role in NDDs since cholesterol and lipids are essential for myelin formation, being myelin alterations important features in many NDDs ^50^. CYP3A4 is localized in the endoplasmic reticulum, and its expression is regulated by glucocorticoids and certain drugs^51^. As we shown in Figure 5, this gene interacts with multiple compounds in the network, five out of eight of which are of plant origin (trifluoroperazine^33^, naringenin^16^, paeonol^18^, apigenin^52^ and quercetin^23^).

The following genes more connected in the gene-compound-disease tripartite network, *CYP1A2*^53^*, CYP2C19, CYP2D6* and *CYP19A1* also encode members of the cytochrome P450 superfamily, like *CYP3A4,* and are part of the same cluster^51^. *CYP2C19* encodes the human liver enzyme CYP2C19, which is also expressed in the human foetal brain. Worth mentioning, humanised CYP2C19 transgenic mice display altered gait and motor impairments, likely resulting from abnormal cerebellar development ^54^. These genes are involved in drug metabolism processes, making it logical that they interact with multiple compounds in the network, regardless of the specific NDDs. This combined interaction with several NDDs is evident in the inferred network (Figure 6) for the genes *CYP3A4, CYP1A2* or *CYP2C19*.

Among the genes with the highest number of interconnections in Figure 5, *ATAD5* differs from the most connected genes in the network (*CYP3A4*, *CYP1A2*, *CYP2C19* and *CYP2D6).* It encodes a protein in humans that plays a crucial role in DNA clamp unloading activity and is involved in the positive regulation of DNA replication and the G2/M phase transition of the cell cycle. Additionally, ATAD5 is the component of the Elg1 RFC-like complex and has been identified as a biomarker for neurilemmoma^55^. Neurilemmoma is a tumour that originates in the peripheral nervous system, specifically from Schwann cells, which are responsible for producing myelin in this system^56^. In our analysis, *ATAD5* interacts with different compounds: quercetin, nimodipine, and entinostat (also known as MS-275). The first is of plant origin, while the latter two are semisynthetic compounds. These three compounds have been studied in the context of Parkinson’s disease (Table 4). According to our reviewed and meta-analysis data, two possibilities emerge. First, that *ATAD5* may be implicated in the Parkinson pathomechanism; and second, that these three compounds, initially explored for Parkinson’s disease, could potentially be repurposed for other conditions, such as neurilemmoma.

Another outstanding gene arose in the tripartite network is *E2F1.* The protein encoded by *E2F1*, which belongs to a family of transcription factors, plays a role in controlling the cell cycle and the action of tumour suppressor proteins. E2F1 is involved in several NDDs such as Alzheimer^57^ and Parkinsońs disease^58^, primarily due to its role in cell cycle regulation, apoptosis, and neuronal cell death.

In line with genes encoding proteins with diverse functions, *CFTR* is highlighted. This gene encodes a member of the ATP-binding cassette (ABC) transporter superfamily. Unlike other members of this family, the encoded protein functions as a chloride channel, playing a crucial role in regulating ion and water transport in epithelial tissues. Its activation is controlled by cycles of regulatory domain phosphorylation, ATP binding to nucleotide-binding domains, and ATP hydrolysis. Mutations in this gene cause adrenoleukodystrophy (ALD), among other diseases^59^. ALD is a fatal progressive NDD affecting brain white matter with limited therapeutic options up to date ^60^, which opens up a venue to expand the potential therapeutic capacity of the reviewed compounds in other NDDs. In the context of this review, compounds interacting with *CFTR* include nimodipine and entinostat (MS-275), as in the case of the *ATAD5* gene, and apigenin. The first two in the context of Parkinson’s disease, while the latter is studied for Hyperglycemia-associated amnesia. This case could be similar to the previous one: a compound studied in the context of one pathology could also be considered for another disease. For instance, Siddaiah et al studied the effect of topiramate, an antiepileptic drug for the treatment of cystic fibrosis^61^. Similarly, apigenin, nimodipine and entinostat could be explored for potential use in other NDDs such as ALD.

In addition, the strong interaction between creatine and the gene *MOBP* is eminently interesting. In humans, *MOBP* encodes myelin-associated oligodendrocyte basic protein (MOBP), which, when is defective, is associated with amyotrophic lateral sclerosis^62^. Amyotrophic lateral sclerosis is a progressive NDD that affects both the upper and lower motor neurons. As shown in Figures 5 and 6, creatine has been explored as a potential drug for the treatment of Huntington’s disease^63^, while our data invites to consider it as a potential treatment for amyotrophic lateral sclerosis as well.

Finally, Huntington’s disease is the most common autosomal dominant neurodegenerative disorder, for which no disease-modifying treatments are currently approved. The molecular pathogenesis of Huntington’s disease is complex, involving toxicity from full length expanded huntingtin (HTT) and its N-terminal fragments, both of which are prone to misfolding due to proteolysis, aberrant intron 1 splicing, and somatic expansion of the CAG repeat in the HTT^64^. However, as verified in the DGIdb database, there is no evidence that the compounds under study for Huntington’s disease directly interact with HTT. Interestingly, we found that terazosin, a compound studied in the context of amyotrophic lateral sclerosis, interacts with HTT^39^. These findings open our perspectives in a double direction, first, that amyotrophic lateral sclerosis and Huntingtońs disease have some common pathomechanism, and that terazosin could be explored further in preclinical models of Huntington’s disease.

## Conclusions

Overall, this review clearly reveals the prominent position of zebrafish as a perfect animal model to develop novel assays in the search of potential high-throughput drug-screening pipelines against NDDs in humans.

NDDs are currently difficult to study, and in many cases, truly effective drugs have yet to be found. This has led to the search for the new compounds that can act on the molecular targets affected in the pathology under study. Given the difficulty in clarifying the causes of the disease and the need to generate animal models that replicate human symptomatology, the use of zebrafish as the preferred model for screening potential drugs is well justified. Due to the involvement of the mitochondrial dysfunction in the pathomechanism of many NDDs, the study of novel compounds with the potential of modulating mitochondrial function stands out as a promising strategy for therapeutic purposes.

This systematic review outlines the broad variety of novel compounds currently studied that can modulate the mitochondrial activity and ameliorate the clinical course of the zebrafish NDD models. In specifics, rodent models, which have also been used historically, sometimes fail to faithfully reproduce disease symptomatology in cases such as Parkinson’s disease. For these reasons, zebrafish is emerging as an alternative, valuable model for screening new potential drugs such as mitochondrial targeted compounds.

Furthermore, the meta-analysis of the reviewed compounds and the genes that these molecules interact with expand the view of the molecular mechanism of action of this new list of drugs, and provides us a more comprehensive picture of the NDD pathologies. To add, the potential of computational tools to predict new therapeutic targets cannot be overlooked. The power of these computational approaches opens the door to generating a series of predictions for drug repurposing or evaluating new compounds even before experimentation begins, potentially saving both time and resources in the research of these diseases.

